# Reference Quality Assembly of the 3.5 Gb genome of *Capsicum annuum* from a Single Linked-Read Library

**DOI:** 10.1101/152777

**Authors:** Amanda M. Hulse-Kemp, Shamoni Maheshwari, Kevin Stoffel, Theresa A. Hill, David Jaffe, Stephen Williams, Neil Weisenfeld, Srividya Ramakrishnan, Vijay Kumar, Preyas Shah, Michael C. Schatz, Deanna M. Church, Allen Van Deynze

**Affiliations:** Department of Plant Sciences, University of California, Davis, CA; USDA-ARS, Genomics and Bioinformatics Research Unit, Raleigh, NC; 10x Genomics, Inc. 7068 Koll Center Parkway, Suite 401, Pleasanton, CA 94566; Department of Computer Science, Johns Hopkins University, Baltimore, MD

**Keywords:** Genome Assembly, *Capsicum annuum*, Complex Genome, Heterozygous, Repetitive Elements, Repeat Content, Linked-Read

## Abstract

**Background:** Linked-Read sequencing technology has recently been employed successfully for *de novo* assembly of multiple human genomes, however the utility of this technology for complex plant genomes is unproven. We evaluated the technology for this purpose by sequencing the 3.5 gigabase (Gb) diploid pepper *(Capsicum annuum)* genome with a single Linked-Read library. Plant genomes, including pepper, are characterized by long, highly similar repetitive sequences. Accordingly, significant effort is used to ensure the sequenced plant is highly homozygous and the resulting assembly is a haploid consensus. With a phased assembly approach, we targeted a heterozygous F_1_ derived from a wide cross to assess the ability to derive both haplotypes for a pungency gene characterized by a large insertion/deletion.

**Results:** The Supernova software generated a highly ordered, more contiguous sequence assembly than all currently available *C. annuum* reference genomes. Eighty-four percent of the final assembly was anchored and oriented using four *de novo* linkage maps. A comparison of the annotation of conserved eukaryotic genes indicated the completeness of assembly. The validity of the phased assembly is further demonstrated with the complete recovery of both 2.5 kb insertion/deletion haplotypes of the *PUN1* locus in the F_1_ sample that represents pungent and non-pungent peppers.

**Conclusions:** The most contiguous pepper genome assembly to date has been generated through this work which demonstrates that Linked-Read library technology provides a rapid tool to assemble *de novo* complex highly repetitive heterozygous plant genomes. This technology can provide an opportunity to cost-effectively develop high-quality reference genome assemblies for other complex plants and compare structural and gene differences through accurate haplotype reconstruction.

## Background

Pursuing a gold-standard reference genome for each biologically important organism has become a goal of the individual research communities in order to have a tool for answering biologically relevant questions [1-5]. The construction of contiguous genome assemblies has allowed for discovery of genes and gene function as well as improved our understanding of genomic elements and structure that regulate biological processes in human, microbes, animals and plants. These high-quality reference genome assemblies allow for not only complete gene models but also complete promoter regions and more remote regulatory sequences of every gene, as well as true representation of other complex features that are important for trait expression. Having a highly accurate, contiguous, and complete genomic representation allows for unprecedented studies on chromosome scale evolution as well as of molecular evolution of polyploid events, gene amplifications, haplotype tracking, and mobile element proliferations. The significance of this completeness and ordering of elements have been continually realized as simple explanations for phenotypes and disease have been lacking, even in human studies with hundreds of thousands of subjects, for quantitative traits such as height where 50-75% of the phenotype has currently been unable to be explained [6, 7].

In plant breeding, the availability of a contiguous genome provides a means to better understand traits and how they interact with their environment in different genetic backgrounds. At the simplest, it allows for association of genetic markers for selection and introgression of traits across germplasm to enable hypothesis-driven crop improvement by understanding the pathways and development of novel products for consumers. A high-quality genome serves as a tool for more efficient studies with higher statistical power for localization of causal genomic regions and genes responsible for economically important traits.

The first high-quality plant genomic sequences were achieved using bacteria artificial chromosome (BAC)-based approaches coupled with Sanger sequencing technology [8], it was limited to a few small diploid species, such as Arabidopsis and rice, due to being labor intensive and expensive with project costs in the tens to hundreds of millions of dollars for a single genome. Next generation sequencing technology, such as Illumina sequencing-by-synthesis, has dramatically reduced costs in the past decade and led to the construction of a large number of draft genomes using a combination of paired-end and mate-pair libraries with short reads and high redundancy. However these draft genomes are usually of low quality and comprised of a large number of contigs (in 100,000s or more) with scaffold N50s in the hundreds of Kilobase pairs (Kb) or less and contig N50s in the tens of Kb, where N50 is the contig or scaffold size at which 50% of the entire assembly is contained. A combination of short read sequencing technology with physical and genetic maps has led to a drastic improvement in scaffold sizes, but not to contigs, leaving many gaps and misassembled or unassembled regions, especially in repeated regions. Long-read technology was introduced in the last decade and lead to dramatic improvements in resolution with N50s over 1 megabase (Mb). However this technology is considerably more expensive than Illumina sequencing (approximately 10 to 20 times more expensive or more) making it difficult to implement with large complex plant genomes. This is particularly problematic for crop genomes which tend to be complex in size, ranging from 0.35 Gb in rice to over 23 Gb in loblolly pine, and are further complicated by polyploidy, varying levels of heterozygosity, and large stretches of highly similar repeat sequence, all of which make their sequence and assembly difficult with standard technologies [9].

To combat the issue of heterozygosity in plants, plant geneticists have taken great concern over choosing a highly homozygous plant for sequencing, such as sequencing highly inbred varieties or by sequencing haploid tissues. This reduced the number of problematic bubbles during computational assembly generation and produced a haploid consensus sequence assembly. However in many species it is not practical to breed homozygous varieties or collect haploid tissue in sufficient quantity for sequencing. Furthermore, generating a haploid consensus sequence will not accurately portray both haplotypes in plants that have varying levels of heterozygous presence/absence variants (PAVs) or insertions/deletions. Some of these PAVs have been shown to be causal for many economically important phenotypes such as pungency in pepper. The *PUN1* gene is a putative acyltransferase in which a 2.5 kb PAV has been shown to be the causal variant in determining a pepper’s distinctive pungent flavor [10]. Having an accurate representation of the PAV regions within an individual line will provide additional power to assess the true biology behind a trait instead of working with a synthetic sequence.

Pepper is a member of the Solanaceae family, which contains several of the most economically important crop species including tomato, potato, eggplant, and tobacco. The pepper genome is a representative complex plant genome; it has one of the largest genome sizes in the Solanaceae family at approximately 3.5 Gigabases and is comprised largely of repetitive elements, estimated at 75-80% of the genome [11, 12]. The most cultivated pepper species (*C. annnum*) is diploid that to-date has three draft genome assemblies developed using short read sequencing technology. All three assemblies focused on a different *C. annuum* line, CM334 which is a Mexican landrace hot pepper [11], Zunla-1 which is a widely cultivated accession and Chiltepin which is a wild progenitor of Zunla-1 [12], each being mostly homozygous due their primary self-pollinating mating type. Similar to most short-read sequencing derived references, the three pepper assemblies are comprised of a large number of small scaffolds with 37,989 scaffolds in CM334 [11], 967,017 scaffolds in Zunla-1 [12], and 1,973,483 scaffolds in the Chiltepin [12] genomes with the largest scaffold N50 at 2.47Mb in the CM334 assembly and largest contig N50 of 55 kb with Zunla. Additional genetic resources have also been recently developed in pepper with one study producing two high-quality manually curated genetic linkage maps with a custom Affymetrix Genechip [13], another study producing a map with an Illumina Infinium array [14], and finally a study producing a map with skim sequencing of a population utilizing the CM334 genome assembly [15]. While multiple short-read assemblies have been implemented in pepper, the large genome size has prohibited sequencing using long-read technology as they would have sequencing costs upwards of $50-100K, currently.

Recent advances in library preparation methods have allowed for integration of structural location information of a sequence with the cost effective short-read sequencing technology. The 10x Chromium technology (10x Genomics, San Francisco, USA) has the potential to make a big impact with complex plant genomes more generally accessible by generating long-range information analogous to old BAC-by-BAC sequencing technologies but at a tiny fraction of the cost and at high throughput. This technology isolates large DNA fragments (^~^150Kb) and creates barcoded Illumina genomic libraries allowing the short reads to be localized by capturing not only sequence, but physical association of the DNA. These libraries are in-turn sequenced to about 60x coverage to use for *de novo* assembly using a phased assembly strategy where individual haplotypes are generated as output [16].

In the current study we have investigated the Linked-Read technology as a cost-effective resource for sequencing the 3.5 Gb complex pepper plant genome, validated the assembly produced by the technology using four high-density genetic maps, and tested the feasibility of the new assembly for answering biologically-relevant questions of the genome structure related to the flavor of the plant with the *PUN1* locus. We show that a significantly improved *de novo* genome assembly can be achieved at a fraction of the time and cost of traditional short-read and long-read assemblies with little input DNA, for a highly-repetitive complex heterozygous pepper plant.

## Results

### Sequencing and Assembly

A total of ^~^56 fold read coverage was obtained with paired-end 150 bp reads using 10x Chromium technology sequenced on the Illumina HiSeq X10 for a single pepper genotype. The sequenced *C. annuum* genotype was an F_1_ wide-cross hybrid of Criollos del Morelos 334 (CM334) and a non-pungent pepper breeding line with a genome wide heterozygosity rate of about 0.4%. The 56X data was assembled using version 1.1d of the Supernova Assembler (https://qithub.com/10XGenomics/supernova-chili-pepper) that utilizes a graph-based assembly approach along with individual molecule barcodes to resolve complex repeats and separate chromosomes based on haplotype information for a phased assembly [16]. The Supernova assembly contained 83,391 scaffold sequences with an N50 of 3.69 Mb for a total assembly size of 3.21 Gb **(Additional File 1).** We validated the overall structure of this assembly by comparing it to four high-density genetic maps available in pepper and found the marker orders in the Supernova assembly was highly concordant to three transcriptome derived maps [13, 14] and one genomic-based map [15] **(Figure 1A, Additional Figures S1-5).** Physical location of markers on the assembly were also compared with the CM334 Pepper Genome V1.55 [11] **(Figure 1B)** which showed that physical location in pericentromeric regions appeared to be more consistent in the 10x assembly positions along the contigs.

**Figure 1.**
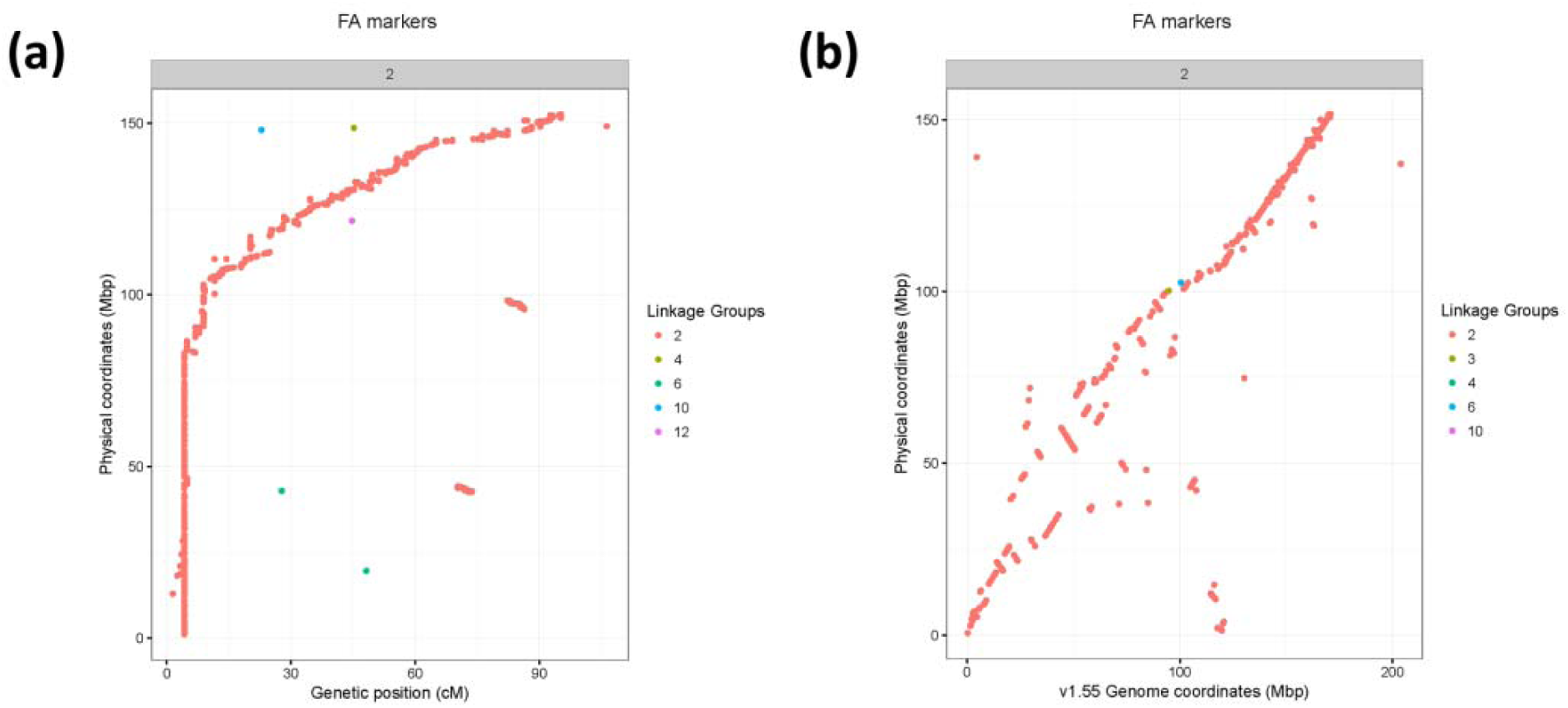
**Assessment of raw 10X assembly contigs** compared to Hill et al. (2015) *Capsicum frutescens* x *Capsicum annuum* genetic map, shown for Chromosome 2. Assembly scaffolds are ordered based on their primary linkage group, sorted in order of increasing genetic distance. Linkage group colored labels correspond to marker linkage group in Hill et al. **A.)** Genetic positions of markers (centiMorgans) are shown versus the physical position on the 10X assembly contigs (megabase pair). **B.)** Physical position on the 10X assembly contigs (Mbp) versus physical position on the C. *annuum* CM334 version 1.55 assembly.

Filtered alignments of marker sequences for the four maps **(Additional File 2)** were utilized in the AllMaps software[17] using a weighted approach to generate pseudomolecules with currently available genetic map information. The highest weights were applied to the highly-manually curated EST maps by Hill et al. [13] and the less manually curated maps were given lower weights [14, 15]. Additionally the intraspecific maps were placed at higher weights while the interspecific maps were given lower weights. This was done to prioritize the within species (*C. annuum*) maps as the interspecific maps had to be corrected for a known translocation in the maps due to the structure of *C. frutescens* compared to *C. annuum* [18]. Chromosome-scale pseudomolecules produced 12 major scaffolds corresponding to the 12 pepper chromosomes **(Figure 2, Table 1, Additional Figures S6-17).** A total of 2.67 Gb was anchored to the 12 chromosomes along with 541 Mb of unplaced sequence for a total assembly of 3.21 Gb designated as UCD10X version 1.0 (available as NCBI bioproject PRJNA376668). The N50 of contigs was 123Kb, of scaffolds was 3.69 Mb, and of pseudomolecules was 227.2 Mb. Over 83% of the assembled sequence was anchored into the final assembly **(Additional File 3).**

**Figure 2.**
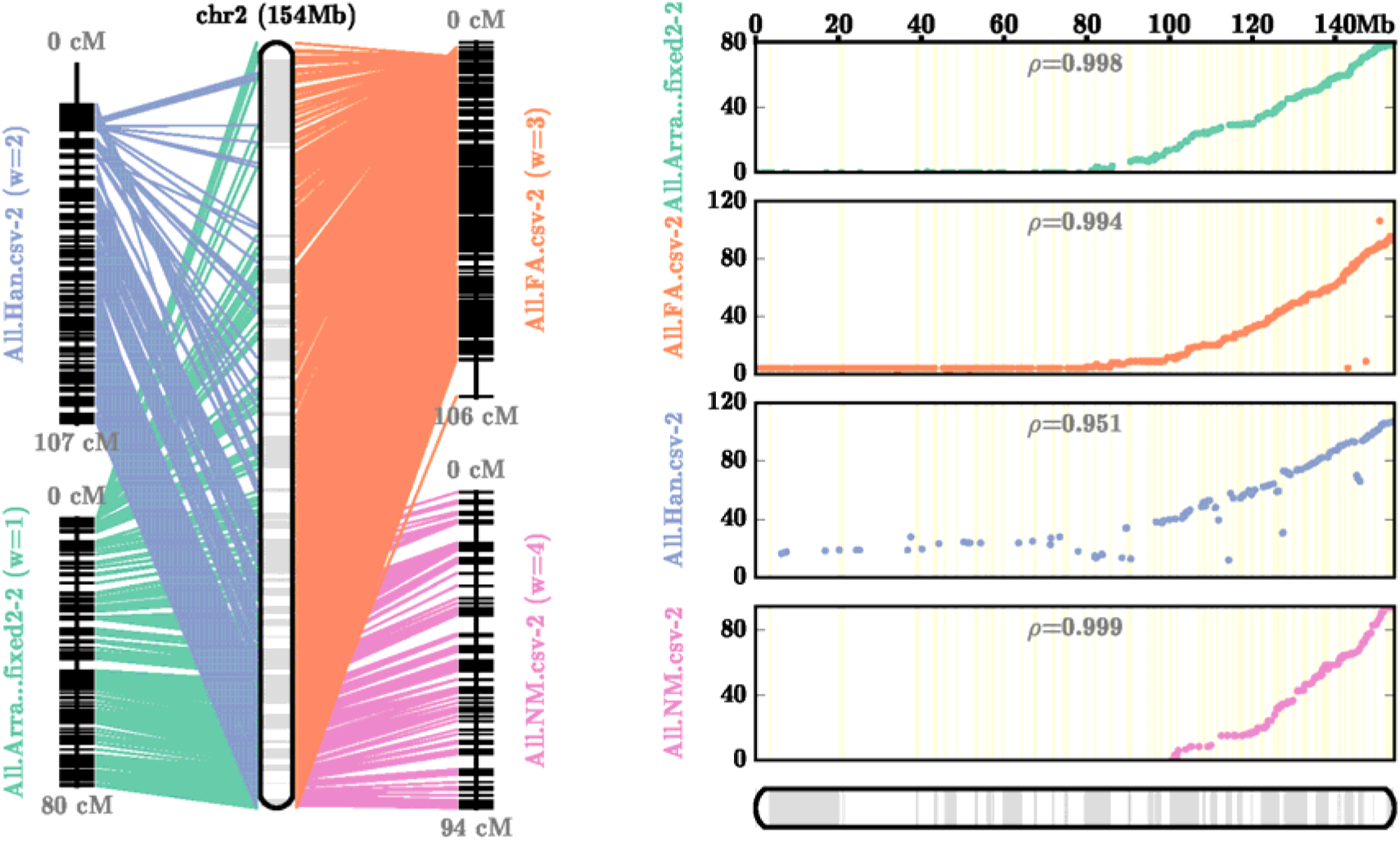
AllMaps Chromosome 2 Consensus Map for Pseudomolecule Generation. Order of markers in the four linkage maps compared to the final pseudomolecule generated through consensus map derivation using the AllMaps software with Unequal Weights2 parameters through whole genome run.

**Table 1.** Chromosome Statistics for UCD10X Genome Assembly.

| Chromosome | Final | Anchored | Oriented |  |  | Gaps | Length |
| --- | --- | --- | --- | --- | --- | --- | --- |
|  | # scaffolds | # scaffolds | # scaffolds | bp | bp % | # N's |  |
| LG01 | 1 | 134 | 51 | 173,703,399 | 67.96% | 13,300 | 255,602,192 |
| LG02 | 1 | 119 | 36 | 79,737,851 | 51.67% | 11,800 | 154,330,250 |
| LG03 | 1 | 153 | 61 | 155,226,247 | 57.37% | 15,200 | 270,589,393 |
| LG04 | 1 | 108 | 44 | 130,088,991 | 56.19% | 10,700 | 231,497,844 |
| LG05 | 1 | 136 | 42 | 121,301,069 | 54.85% | 13,500 | 221,136,838 |
| LG06 | 1 | 136 | 39 | 99,428,383 | 43.56% | 13,500 | 228,254,876 |
| LG07 | 1 | 139 | 40 | 81,805,368 | 36.01% | 13,800 | 227,195,441 |
| LG08 | 1 | 61 | 25 | 127,432,512 | 73.33% | 6,000 | 173,776,113 |
| LG09 | 1 | 203 | 28 | 75,652,043 | 34.53% | 20,200 | 219,064,469 |
| LG10 | 1 | 121 | 43 | 116,352,171 | 52.48% | 12,000 | 221,721,387 |
| LG11 | 1 | 143 | 35 | 106,810,631 | 45.27% | 14,200 | 235,950,708 |
| LG12 | 1 | 134 | 43 | 122,946,399 | 52.77% | 13,300 | 232,995,803 |
| **Total** | **12** | **1,587** | **487** | **1,390,485,064** | **52.04%** | **157,500** | **2,672,115,314** |
| Unplaced | 81,804 | - | - | - | - | - | 541,059,453 |
| **Including Unplaced** | **81,816** | **1,587** | **487** | **1,390,485,064** | **43.27%** | **157,500** | **3,213,174,767** |

### Comparison to Published Sequences

UCD10X was compared to the three other publicly available *C. annum* genome sequences: (1) CM334 version 1.55 [11], (2) Zunla-1 version 2.0, and (3) Chiltepin version 2.0 [12] using QUAST [19]. The GC% of UCD10X was 34.91%, comparable to the other published assemblies, which ranged from 34.97% to 35.09%. The length of sequence anchored to pseudochromosome scaffolds ranked second among the assemblies at 2.67 Gb anchored, compared to CM334 (2.75), Zunla-1 (2.65) and Chiltepin (2.45). The overall size of the assembly (3.21 Gb) was also within the range of the other assemblies.

While the assemblies at the overall level of pseudochromosomes appeared comparable, the quality within the pseudochromosomes was variable, especially when the assemblies were resolved to their constituent contigs. The UCD10X assembly contained the smallest number of contigs (134,573), with the next most contiguous genome having 32% more contigs (CM334 v1.55 - 177,870). The UCD10X assembly also has a contig N50 of 123kb, 2x greater than the other three genomes. Ultimately the UCD10X produced the most contiguous assembly with over 75% of the total sequence length in contigs over 50Kb **(Figure 3).**

**Figure 3.**
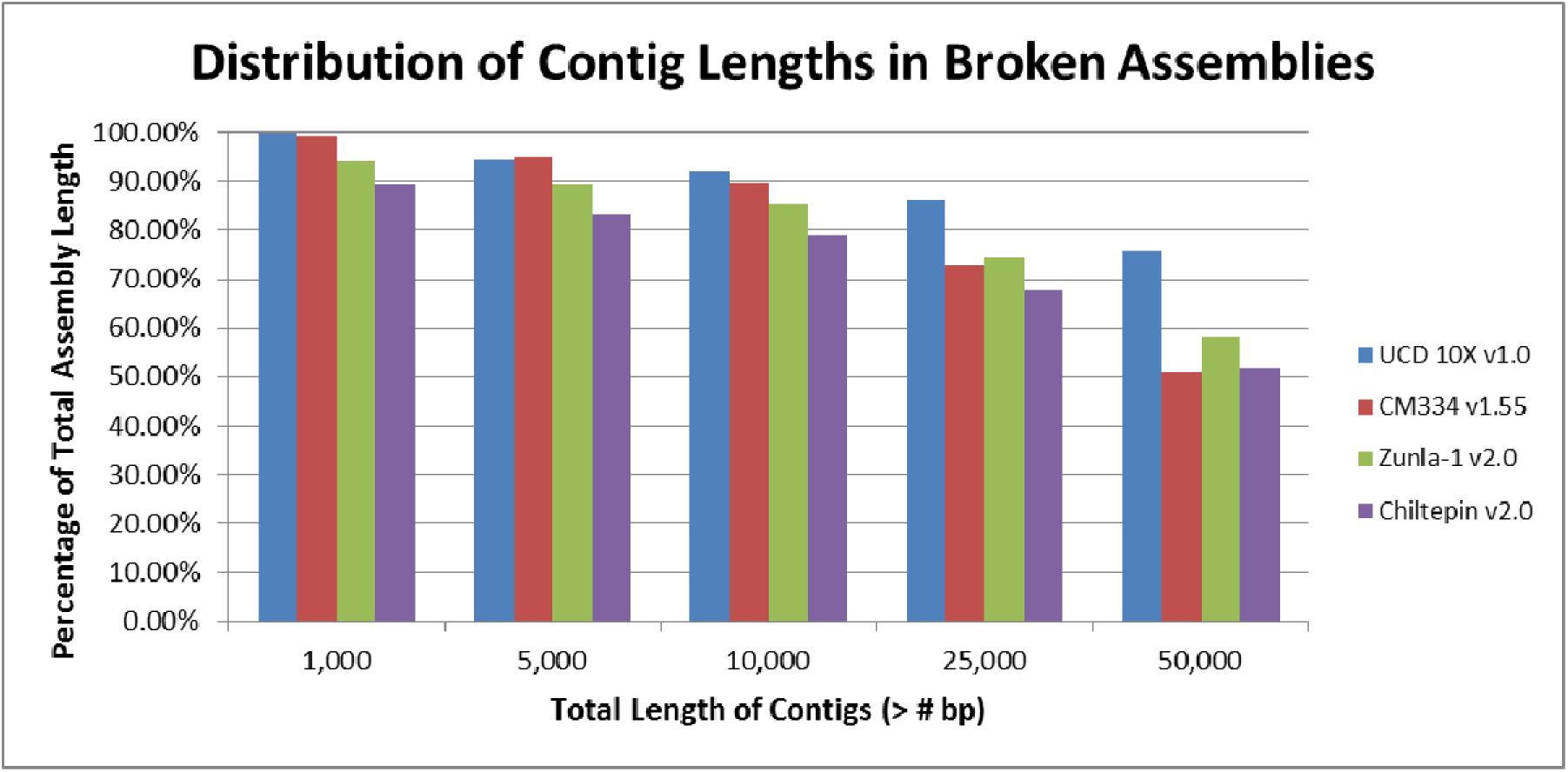
Distribution of Contig Lengths in Broken Assemblies. The proportion of the total assembly length found in contigs that are over a range of base pairs in the four *Capsicum annuum* published genome sequences.

Gene content of UCD10X, the three other Capsicum genomes, as well as assembly sequences for other Solanaceae (Tomato [20], Potato [21], Tobacco [22], Eggplant [23], and Petunia [24]) and an asterid outgroup (Carrot [25]) were assessed using BUSCO2 [26], a standard benchmarking software for assessing genome completeness by measuring the number of core genes present and full length in the assemblies. The embryophyta_0db9* standard data set includes 1,440 genes that are conserved among 90 representative plants. All runs utilized tomato as the training species for gene model detection and UCD10X was found to perform comparably to the other pepper genomes containing 1,336 (92.78%) of the total genes in complete copies **(Figure 4).** The diploid Solanaceae including Potato, Eggplant, and Pepper appeared to all have similar numbers of fully duplicated conserved genes at 38-39, while Petunia had closer to the Carrot outgroup and polyploid Tobacco had high numbers of duplicated genes as expected.

**Figure 4.**
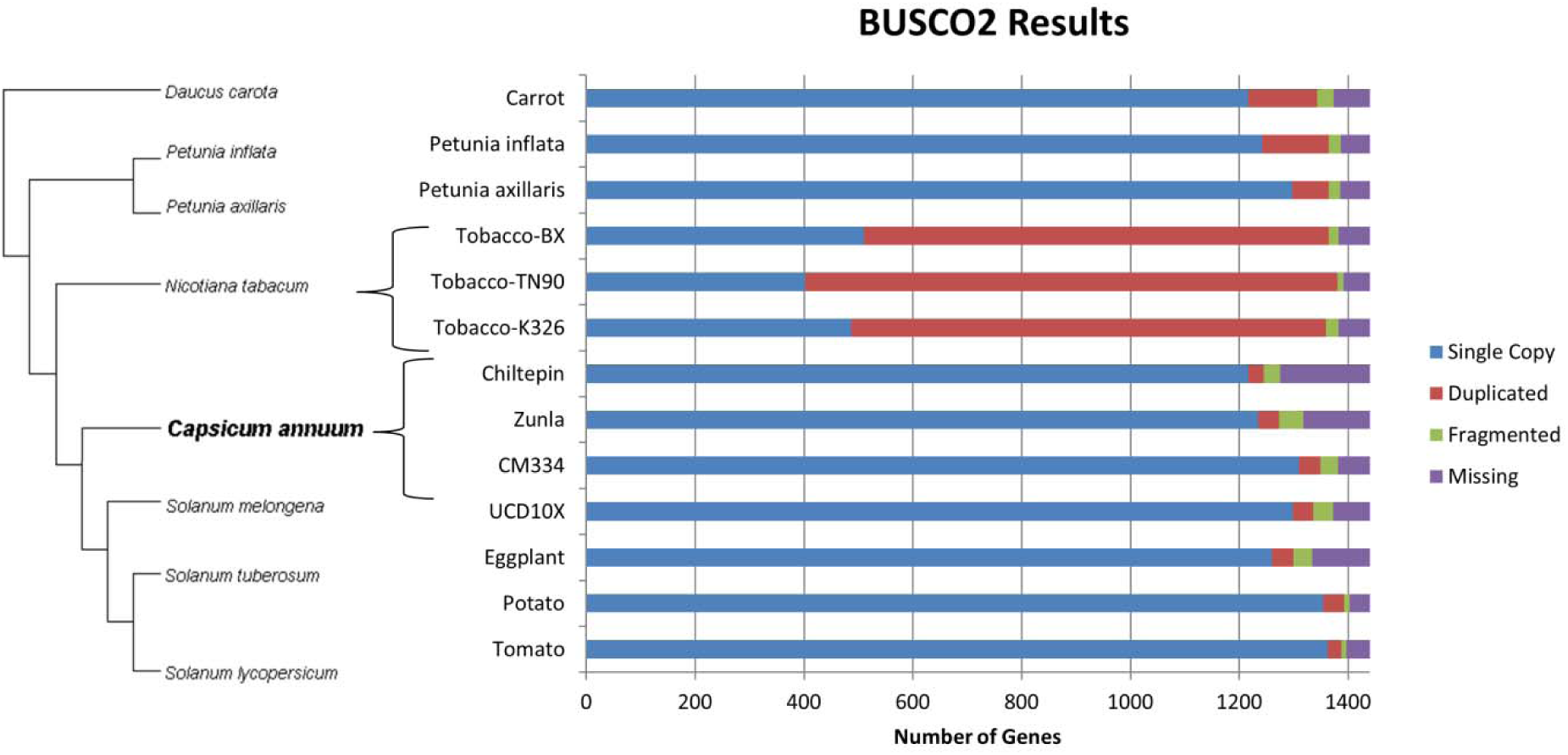
BUSCO2 Conserved Gene Analysis Comparison. BUSCO2 results for the UCD10X assembly, three other pepper assemblies, other available Solanaceae family genomes including Tomato, Potato, Tobacco, Eggplant and Petunia, and Carrot as an asteroid outgroup.

The final pseudomolecules showed high congruence with the other pepper genomes over the euchromatic, non-centromeric regions of the chromosomes, as shown on the long arm of Chromosome 2 **(Figure 5, Additional Figures S18-21).** Chromosome 2 is an acrocentric chromosome with a very small short arm compared to the long arm [27]. The heterochromatic region appears to be highly variable between all of the assemblies, except for the comparison between Zunla-1 and Chiltepin. A similar pattern of higher collinearity in euchromatic regions and lower collinearity in centromeric regions is observed for all pepper assemblies when comparing chromosome 2 to the orthologous chromosome 2 in tomato **(Figure 6).** Furthermore when the syntenic region is examined closer, the alignment of the UCD10X sequence appears to be more contiguous and contain better oriented contigs overall compared to tomato with less switching between plus (red) and minus (blue) orientation in the comparison. As tomato is a closely related Solanaceae member, it is expected that the sequences would show high co-linearlity. Determining the correct sequence orders between related species is desired to allow for identification of high-resolution syntenic relationships and assessment of gene orthology for comparative studies [28].

**Figure 5.**
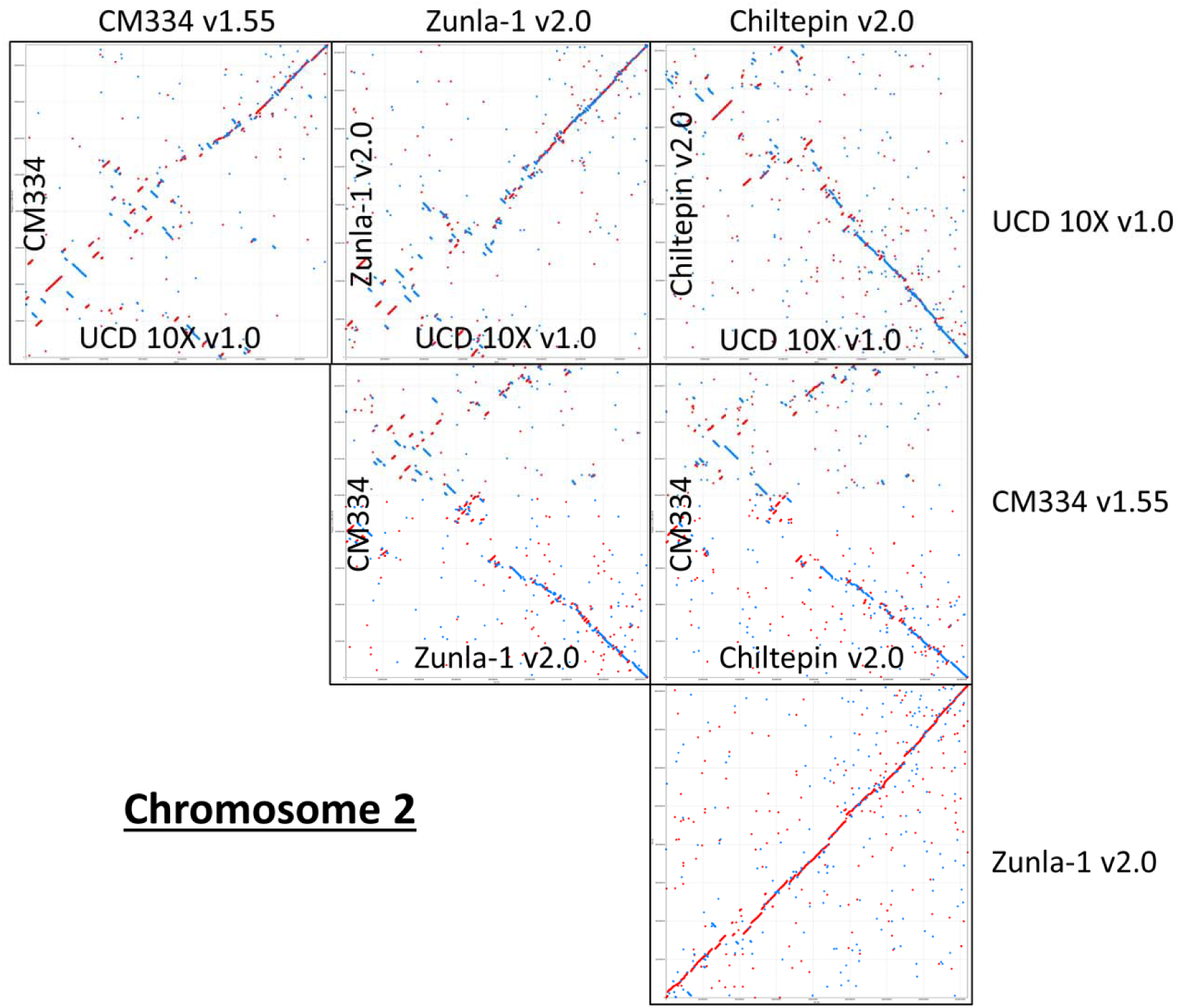
Comparison of 10x Assembly versus other Pepper assemblies. Filtered nucmer alignments extracted for Chromosome 2 are shown for alignment lengths of greater than 150 base pairs at greater than 95% sequence identity. Pairwise alignments between the UCD10X version 1.0, CM334 version 1.55, Zunla version 2.0 and Chiltepin version 2.0.

**Figure 6.**
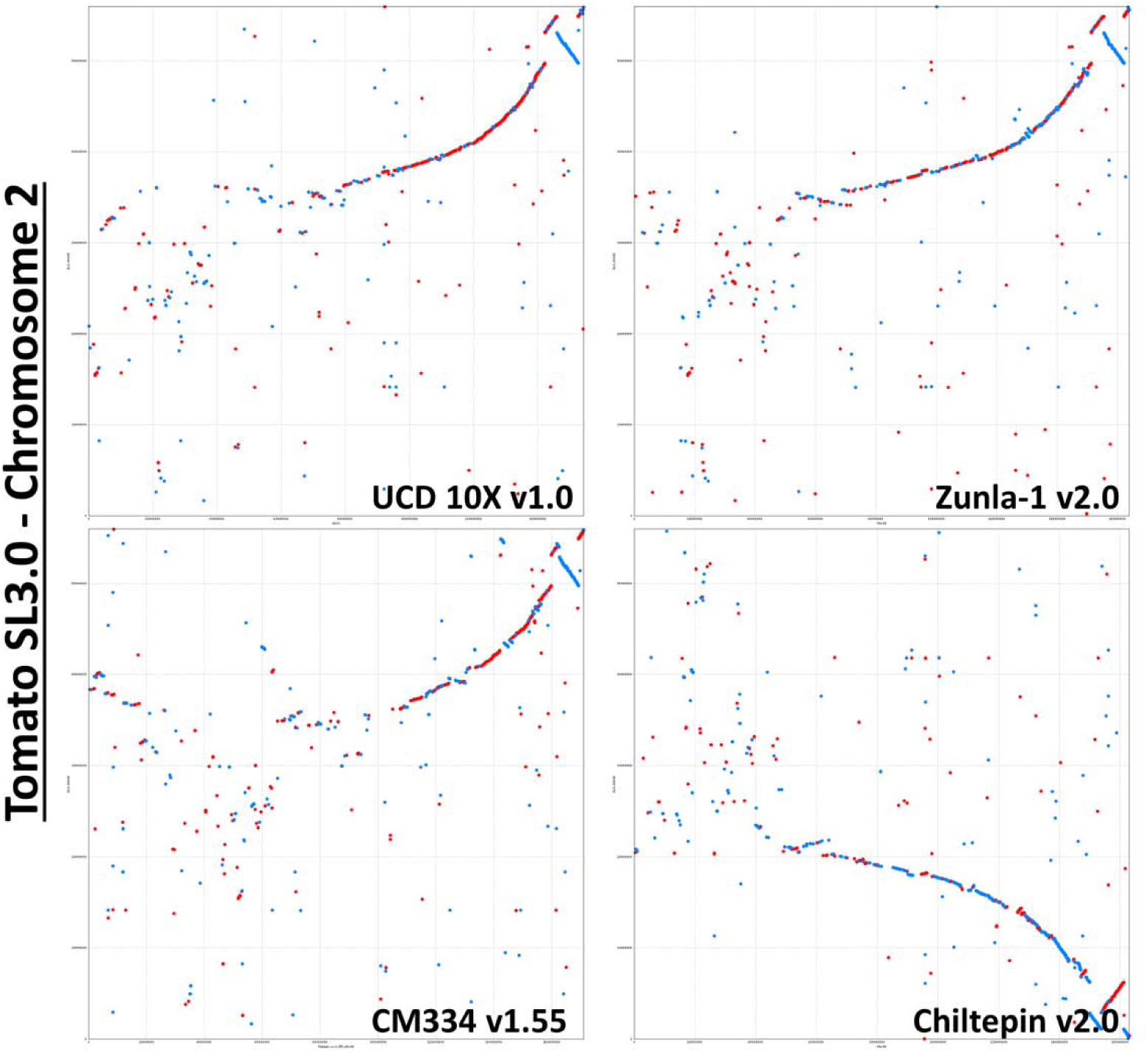
Comparison of 10X Assembly and other Pepper assemblies against Tomato. Filtered nucmer alignments extracted pepper Chromosome 2s and *Solanum lycopersicum* (Tomato version SL3.0) are shown for alignment lengths of greater than 150 base pairs at greater than 85% sequence identity. Pairwise alignments with tomato are shown with UCD10X version 1.0, CM334 version 1.55, Zunla version 2.0 and Chiltepin version 2.0 on the X-axis and Tomato on the Y-axis.

We further analyzed the four pepper assemblies reference contigs with the whole genome MUMmer [29] alignments with Assemblytics [30] to discover any structural variant that ranges within 1 to 10,000 base pairs. This approach depends on having a high quality assembly, as it is not possible to find large structural variations if the assembly is highly fragmented. Assemblytics reported that the UCD10X showed the most number of structural variations and most total bases affected by structural variants of all the four pepper assemblies **(Table 2).** It is also reported that UCD10X captured the most repeat expansions with 6% more events, which covers 20% more in terms of total bases than other pepper assemblies. While there are undoubtedly some genuine biological differences between the four pepper varieties, we attribute the increased sensitivity of structural variants in the UCD10X assembly to its superior quality. This also suggests that the other three assemblies may be missing the sequences for many thousands of variants.

**Table 2.** Comparison of variant counts between Pepper Genome Assemblies and Tomato Reference

| Assemblytics Comparison of Variant Counts Between Pepper Genome Assemblies and Tomato Reference |  |  |  |  |  |  |  |  |
| --- | --- | --- | --- | --- | --- | --- | --- | --- |
|  | C. annum UCD10X v1.0 |  | C. annum Zunla v2.0 |  | C. annum CM334 v1.55 |  | C. annum Chiltepin v2.0 |  |
|  | Count | Total bp | Count | Total bp | Count | Total bp | Count | Total bp |
| **Insertion** |  |  |  |  |  |  |  |  |
| 1-10bp | 0 | 0 | 0 | 0 | 268,433 | 517,129 | 240,128 | 463,867 |
| 10-50bp | 3 | 83 | 1 | 27 | 5,809 | 84,723 | 5,127 | 73,749 |
| 50-500bp | 791 | 166,658 | 776 | 167,354 | 852 | 167,852 | 781 | 160,426 |
| 500-10,000bp | 297 | 521,828 | 283 | 488,244 | 295 | 537,853 | 245 | 4,133,314 |
| Total | 1,091 | 688,569 | 1,060 | 655,625 | 275,389 | 1,307,557 | 246,281 | 1,111,356 |
| **Deletion** |  |  |  |  |  |  |  |  |
| 1-10bp | 0 | 0 | 0 | 0 | 270,354 | 501,192 | 239,699 | 445,151 |
| 10-50bp | 0 | 0 | 0 | 0 | 4,752 | 68,332 | 4,150 | 59,493 |
| 50-500bp | 503 | 106,777 | 493 | 103,119 | 535 | 108,600 | 492 | 100,568 |
| 500-10,000bp | 186 | 269,673 | 190 | 268,250 | 190 | 282,310 | 162 | 219,217 |
| Total | 689 | 376,450 | 683 | 371,369 | 275,831 | 960,434 | 244,503 | 824,429 |
| **Tandem_expansion** |  |  |  |  |  |  |  |  |
| 1-10bp | 0 | 0 | 0 | 0 | 0 | 0 | 0 | 0 |
| 10-50bp | 0 | 0 | 0 | 0 | 0 | 0 | 0 | 0 |
| 50-500bp | 21 | 3,531 | 53 | 13,227 | 43 | 8,931 | 55 | 13,695 |
| 500-10,000bp | 164 | 794,086 | 434 | 1,499,840 | 250 | 1,117,619 | 350 | 1,134,864 |
| Total | 185 | 797,617 | 487 | 1,513,067 | 293 | 1,126,550 | 405 | 1,148,559 |
| **Repeat_expansion** |  |  |  |  |  |  |  |  |
| 1-10bp | 264 | 1,312 | 262 | 1,329 | 260 | 1,307 | 226 | 1,179 |
| 10-50bp | 1,107 | 33,406 | 1,105 | 33,012 | 1,063 | 31,942 | 1,019 | 30,214 |
| 50-500bp | 5,824 | 1,283,246 | 56,778 | 1,252,312 | 5,445 | 1,184,547 | 5,417 | 1,195,192 |
| 500-10,000bp | 7,168 | 19,008,122 | 6,611 | 16,529,538 | 5,127 | 11,420,087 | 6,082 | 14,816,116 |
| Total | 14,363 | 20,326,086 | 13,656 | 17,816,191 | 11,895 | 12,637,883 | 12,744 | 16,042,701 |
| **Repeat_contraction** |  |  |  |  |  |  |  |  |
| 1-10bp | 236 | 1,194 | 243 | 1,211 | 242 | 1,199 | 233 | 1,136 |
| 10-50bp | 954 | 27,646 | 983 | 28,908 | 938 | 27,233 | 857 | 24,819 |
| 50-500bp | 3,547 | 742,313 | 3,429 | 732,281 | 3,391 | 713,492 | 3,219 | 674,573 |
| 500-10,000bp | 2,414 | 4,670,668 | 2,335 | 4,483,897 | 2,138 | 3,901,448 | 2,110 | 3,968,958 |
| Total | 7,151 | 5,441,821 | 7,020 | 5,246,297 | 6,709 | 4,643,372 | 6,419 | 4,669,486 |

### Accuracy of Haplotype Reconstruction

All available full-length *PUN1* gene sequences from NCBI were obtained and aligned to the UCD10X assembly to determine the position of the gene in the assembly. The gene was found to be located on chromosome 2 over positions 135,884,368 to 135,885,734. This region and the corresponding region in the alternative haplotype were aligned along with the gene sequences obtained from NCBI using MUSCLE software [31]. Sequence alignments showed two distinct sequence types for pungent and nonpungent lines, highlighting the importance of a phased diploid genome assembly **(Figure 7).** Specifically, the haplotype sequence from contig 3924 in UCD10X clustered with the genes derived from nonpungent *C. annuum* lines while the corresponding haplotype sequence from contig 3922 clustered with genes derived from the pungent lines sequenced individually, indicating the complete reconstruction of the 2.57 kb hemizygous deletion of the *PUN1* gene haplotypes in the sequenced individual.

**Figure 7.**
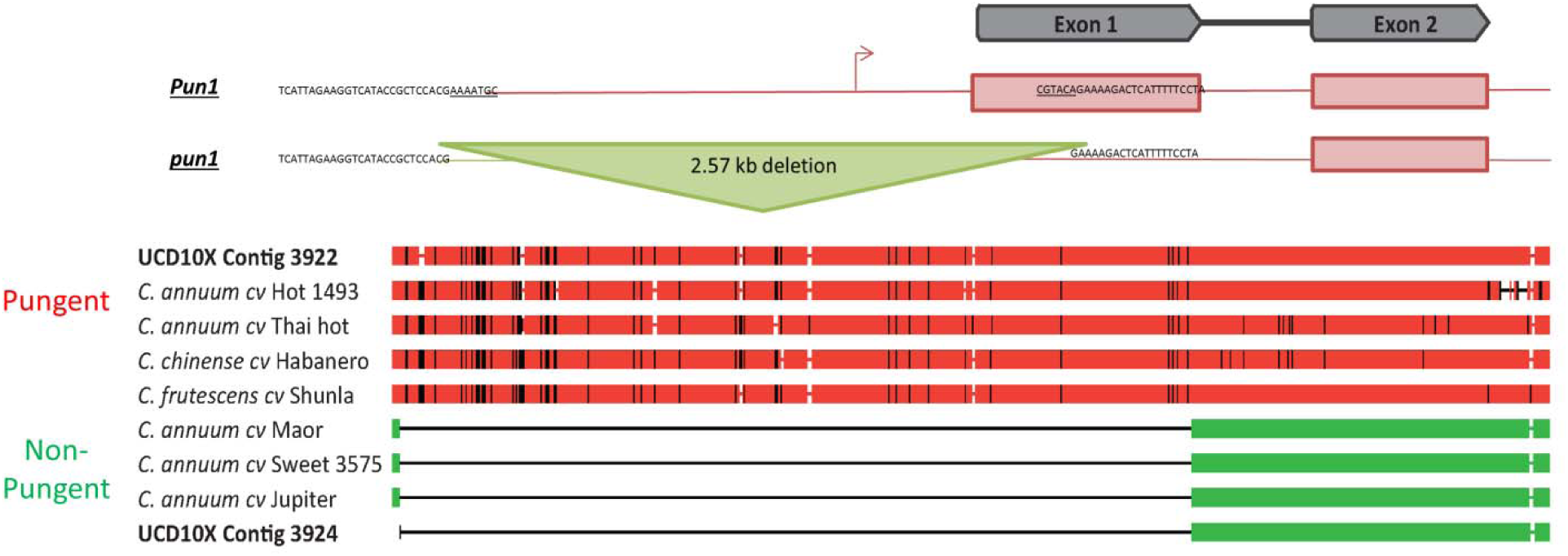
Analysis of Pun1 gene assembly sequence. Structure of Pun1 locus and the corresponding haplotypes of 7 Pun1 gene sequences obtained from NCBI and the loci extracted from both UCD10X assembly haplotypes. Full size of deletion in the alignments is 2,574 base pairs. Sequence alignments show high sequence similarity and separation of pungent and non-pungent groups.

## Discussion

Our newly assembled pepper genome sequence assembly, UCD10X, was found to be of the highest contiguity of any published pepper genome despite being produced with an F_1_ individual. This test case has shown that it is possible with the 10x Chromium Linked-Read technology to accurately assemble and recover both parental haplotype sequences while sequencing a single individual, as demonstrated in the *PUN1* region. Recovery of both haplotypes with 10x Chromium Linked-Reads is a powerful advance over standard short-read sequencing, in that short reads inherently provide low power for discovery and *de novo* reconstruction of genomes, especially in large heterozygous samples. Furthermore, these benefits are generated at a very reasonable cost of data acquisition (^~^$6K for 3.5Gb genome), and at a fraction of the costs for traditional short read strategies and long-read sequencing. This opens the exciting possibility to make linked read sequencing and *de novo* assembly a routine operation, on par with regular sequencing efforts, but with much greater power to detect structural variations and haplotype differences.

While the current assembly is highly contiguous and has resolved the haplotypes over much of the genome, some challenges remain in the sequencing and assembly of complex plant genomes and some regions may not be able to be accurately assembled with Linked-Reads. Large portions of complex plant genomes are comprised of repetitive elements. Some of the longest repeats may span distances longer than the individual molecule lengths, which will still cause breaks in contigs across these regions. Consequently, the success with 10x Chromium linked-reads or alternative long read sequencing systems will be dependent on the distribution and length of the repetitive elements in the genome. The repetitive content in pepper constitutes >80% of the genome and was derived by a rapid expansion of retrotransposon elements (70% of the genome), mainly of a single Gypsy element family, after divergence from the other Solanaceae members [11, 12]. While these repetitive elements may have caused fragmentation of the genome, the overall product was very contiguous with over 50% of the genome in the *de novo* assembled scaffolds larger than 3.69Mb. This allowed for most (83%) of the total assembly length to be placed into pseudomolecules using the four available *Capsicum* linkage maps. Only very small scaffolds could not be confidently placed in the final pseudomolecules as expected **(Additional Figure S22).**

The chromosome sizes determined for UCD10X through use of the four linkage maps are also comparable to the other published genome assemblies **(Additional Figure S23).** The notable differences are with Chromosomes 1 and 8, which are known to have a translocation between *C. annuum* and *C. frutescens.* This is important as these species were used to generate populations with a high-polymorphism rate for genetic mapping [13, 14]. The breakpoint in the two interspecific maps in this particular case was pinpointed with manual hand-annotation by Hill et al. (2015) to ensure that genetic map data was correctly associated with the corresponding *C. annuum* chromosome. It can be seen that this caused UCD10X Chromosome 1 to be slightly smaller than the other three assemblies while UCD10X Chromosome 8 is larger than the other assemblies and closer in size to the other *C. annuum* chromosomes which would be expected based on the pepper karyotype where most chromosomes are similar in size [32].

Linked-Read genomic library technology paired with short read sequencing have made it possible to generate long contiguous scaffolds for pepper and moderately-sized contigs that was previously not possible through short-read sequencing and considerably less expensive than would be needed for long-read sequencing. This experiment has shown that it is possible to sequence large complex plant genomes such as pepper using the 10x Chromium technology and customized, open source Supernova assembler. These tools will make it possible to broaden the scope of high-quality draft assemblies in an economically feasible manner for crops which are limited in funding. It also makes it possible to sequence large collections of individual genomes to very high-quality, something not tractable with more expensive long-read sequencing. A similar strategy will make molecular breeding tools more accessible for more crops and advancements to be made at a quicker pace to assist in providing nutritious food for a growing world population.

## Conclusions

A highly contiguous assembly for a heterozygous complex genome of pepper has been generated in an economically viable manner through Linked-Read sequencing technology that combines the cost efficiency of short-read sequencing with barcoded sequencing libraries that retain long-range physical information. Importantly, the technology allowed for contiguity across long repeats and pericentromeric regions. We showed that large heterozygous (hemizygous) structural variants can be defined in a single *de novo* assembly, which provides an opportunity to cost-effectively compare structural and gene differences among *de novo* sequence assemblies among genotypes rather than simply mapping reads to a reference genome. This technology changes the type of research attainable for plant breeding; functional analyses of genes and genomic elements; and our understanding of genome evolution across complex organisms.

## Methods

### Plant material and DNA extraction

High molecular weight DNA was isolated using a modified CTAB protocol [33] in a F_1_ of a wide cross (UCD-10X-F_1_) between a landrace, Criollos del Morelos 334 (CM334) and a non-pungent blocky pepper breeding line. The only protocol modification was running the Pippen Pulse gel on the 5-150 Kb setting to perform size selection of fragments over 48kb **(Additional Figure S4).**

### Library construction and sequencing

High molecular weight DNA (1.25 ng) was loaded onto a Chromium controller chip, along with 10x Chromium reagents and gel beads following manufacturers recommended protocols (https://support.10xqenomics.com/de-novo-assemblv/librarv-prep/doc/user-quide-chromium-qenome-reaqent-kit-v1-chemistrv). Briefly, initial library construction takes place within droplets containing beads with unique barcodes (called GEMs). The library construction incorporates a unique barcode that is adjacent to read one. All molecules within a GEM get tagged with the same barcode, but because of the limiting dilution of the genome (roughly 300 haploid genome equivalents) the probability that two molecules from the same region of the genome are partitioned in the same GEM is very small. Thus, the barcodes can be used to statistically associate short reads with their source long molecule. The resulting library was sequenced on two lanes of an Illumina HiSeq X10 sequencer to produce 2x150 paired-end sequences. The resulting data type is called ‘Linked-Reads’ [34]. Raw data has been uploaded to NCBI’s Small Read Archive (will be added upon publication SRS#####).

### Assembly & Assembly Verification

The Linked-Read data was assembled with version 1.1d of the Supernova™ assembler [16] using the default recommended settings available on GitHub: https://qithub.com/10XGenomics/supernova-chili-pepper. Molecule size following sequencing of the Chromium library was estimated using the LongRanger tool from 10x Genomics **(Additional Figure S25).** To verify the quality of the assembly we compared the order of contigs to four high density inter- (*C. annuum)* and intra-specific (*C. annuum* x *C. frutescens*) genetic maps: three transcriptome based maps [13, 14] and one genomic map [15]. Marker and/or flanking sequences for all maps were aligned as query sequences to the Supernova 10x assembly using GMap v09-14-2016 with default settings [35].

Alignment results were filtered for hits obtaining >90% of the query sequence of at least 200 base pairs aligning to the target at 98% sequence identity. In the case of the Array map [14], the cut off for size of query sequence was >100 bp to conform to the assayed sequences on the array. The number of scaffolds that a marker aligned to were examined; after filtering, the majority of the markers were homologous to a single scaffold **(Additional Figures S3A-D).** For further analyses, markers/ESTs that aligned to more than one 10x scaffold were removed. Alignment positions (Mb) of the linkage mapped markers were plotted against their centiMorgan (cM) position in the linkage map **(Figure 1A and Additional Figures S4-7A),** if markers/ESTs from >1 linkage group aligned to the same 10x scaffold then the primary linkage group of that scaffold plotted is that of the majority of corresponding markers. The 10x scaffolds belonging to the same primary linkage group were sorted in order of increasing genetic distance of their aligned ordered markers. Additionally, where previously reported positions of markers on the Kim et al. [11] assembly were available, their positions were plotted against aligned position on the 10x scaffolds **(Additional Figures S4-7B),** where the 10x scaffolds were first assigned primary chromosomes using similar logic previously mentioned and then sorted in order of the location of the marker on the Kim et al. genome sequence.

Marker alignment positions on the 10x scaffolds were converted into csv format for input into AllMaps software [17] for all four linkage maps **(Additional File 2).** Interspecific linkage map files were modified to correct for a known translocation between chromosomes 1/8 between *C. annuum* and *C. frutescens* [18]. AllMaps was initially run for sets of markers representing a single chromosome with five different parameter sets for the array map [14], FA map [13], Han map [15], and NM map [13] as follows: 1) Equal weights, 2) Unequal Weights1-1/2/2/2, 3) Unequal weights2 – 1/3/2/4, 4) Unequal weights3 – 1/3/2/3, and 5) Unequal Weights4 – 2/3/1/4. The best result of the five parameter set was determined by greatest number of anchored and oriented scaffolds.

### Pseudomolecule Construction

Final pseudomolecules were constructed in AllMaps using the unequal weights2 parameters for a single AllMaps run for the entire genome. The resulting final pseudomolecules were deposited to NCBI under BioProject ID PRJNA376668, designated UCD10X Assembly v1.0. Assembly statistics through each step of the analysis from contigs to scaffolds to pseudomolecules were calculated.

### Comparison of assembly with published assemblies

Quast [19] (version 4.1) was utilized to simultaneously compare the UCD 10X Assembly v1.0 with three published *C. annuum* sequences, CM334 v1.55 [11], Zunla-1 v2.0 [12] and Chiltepen v2.0 [12]. Default parameters were used except for a minimum contig size of 45 bp and using the scaffold option to also compare broken assemblies. Distributions of contig lengths of the broken assemblies were plotted.

All four pepper assemblies and the *Solanum lycopersicum* version 3.0 (Tomato) were aligned against each other in a pairwise fashion for all pseudochromosome sequences using Mummer version 3.23 [29]. The nucmer alignment algorithm was utilized requiring minimum clusters of 100 base pairs with maximum gap of 500 base pairs. These results were then filtered for the within pepper comparisons for alignment lengths of >500 bp at 98% identity and for the pepper to tomato comparisons for alignment lengths of >150bp at 85% identity. These filtered alignments were plotted for visualization using the mummerplot function **(Additional File: Figures S8-15).** Chromosome 2 alignments were extracted from the overall alignments as a highlight in **Figures 5** & **6.** Nucmer –maxmatch algorithm results with minimum alignment length set to 20 base pairs and minimum length clusters of 100 base pairs were extracted from the four pepper assemblies aligned to the Tomato and analyzed with Assemblytics software [30].

The benchmarking software BUSCO version 2.0[26] was run against all four pepper assemblies (UCD10X, CM334, Zunla, Chiltepin), Tomato – *S. lycopersicum* version 3.00, Potato – *S. tuberosum* version 3, Eggplant – *S. melongeno* version 2.5.1, three Tobacco – *Nicotiana tobocum* genomes (K326, TN90, BX), two *Petunia* genomes (*axillaris, inflata*) and Carrot – *Daucus carota* version 2.0 as a related outgroup with dependencies of BLAST 2.6.0, HMMER 3.1b2, and AUGUSTUS 3.2.3 [26]. The all plant ancestry set, embryophyta_odb9, was used as a reference and all runs utilized tomato species parameters through the ‘-sp tomato’ option. The number of complete single-copy, complete duplicated, fragmented and missing BUSCOs were calculated and compared **(Figure 3).**

### Assessment of *PUN1*

All *PUN1 Capsicum* full-length coding sequences were obtained from the NCBI database (AY819027.1, AY819026.1, AY819029.1, EF104910.1, HM854860.1, GU300812.1, AY819032.1, AY819031.1, AY819030.1). All sequences were aligned with default BLAST alignment parameters to the UCD10X v1.0 pseudomolecules to identify the PUN1 sequence. The identified region and the corresponding region on the other haplotype sequence were extracted. A multiple sequence alignment was generated using MUSCLE software for both extracted sequences and the 7 full-length PUN1 sequences from NCBI. The multiple sequence alignment was analyzed in JalView 2.10.1 and the first three bases of haplotype 2.1 was corrected to positions 24-26. The manually curated alignment was exported in FNA format and visualized **(Figure 3).**

## List of Abbreviations

Gb: Gigabase
BAC: Bacteria artificial chromosome
Kb: Kilobase
Mb: Megabase
PAV: Presence absence variant
CM334: Criollos del Morelos 334
QTL: Quantitative trait loci
cM: CentiMorgan

## Acknowledgements

The authors would like to thank the University of California, Davis Genome Center for maintenance and support of the computational resources utilized for this project.

## Declarations

### Availability of data and material

The datasets generated and analyzed during the current study are available in the NCBI database under BioProject ID PRJNA376668. All other data generated or analyzed during this study are included in this published article and its supplementary information files.

### Authors’ contributions

AVD designed and conceived the project. AMHK, SM, TAH, and DMC designed and performed the analyses. KS, DJ, SW, NW, SR, VK, PS, SR, and MCS performed the analyses. AMHK wrote the manuscript. All authors read, edited, and approved the manuscript.

## List of Additional Files

**Additional File 1** – UCD10X Assembly Statistics.

**Additional File** 2 – Marker Locations on UCD10X contigs used to generate pseudomolecules through four genetic linkage maps

**Additional File** 3 – Chain file of Psuedomolecule Generation from Scaffolds.

